# Cold induces brain region-selective cell activity-dependent lipid metabolism

**DOI:** 10.1101/2024.04.15.589506

**Authors:** Hyeonyoung Min, Yale Y Yang, Yunlei Yang

## Abstract

It has been well documented that cold is an enhancer of lipid metabolism in peripheral tissues, yet its effect on central nervous system lipid dynamics is underexplored. It is well recognized that cold acclimations enhance adipocyte functions, including white adipose tissue (WAT) lipid lipolysis and beiging, and brown adipose tissue (BAT) thermogenesis in mammals. However, it remains unclear whether and how lipid metabolism in the brain is also under the control of cold acclimations. Here, we show that cold exposure predominantly increases the expressions of the lipid lipolysis genes and proteins within the paraventricular nucleus of the hypothalamus (PVH). Mechanistically, we find that cold activates cells within the PVH and pharmacological inactivation of cells blunts cold-induced effects on lipid peroxidation, accumulation of lipid droplets (LDs), and lipolysis in the PVH. Together, these findings suggest that PVH lipid metabolism is cold sensitive and integral to cold-induced broader regulatory responses.

## Introduction

Lipid metabolism in peripheral tissues has been extensively studied, including white adipose tissue (WAT) lipolysis (*Grabner et al., 2021*), beiging (*Bartelt and Heeren, 2014*) and brown adipose tissue (BAT) thermogenesis (*Carpentier et al., 2023*); however, an essential but poorly understood element is that of lipids in the central nervous system, particularly in the hypothalamus which plays a crucial role in the regulation of systematic energy metabolism (*Waterson and Horvath, 2015*) and glucose homeostasis (*Pozo and Claret, 2018*). The brain is highly thermal sensitive in that one °C or less temperature changes can lead to functional alterations of the central nervous system (*Brooks, 1983; Wang et al., 2014*). Lipids provide a major source of energy and heat in the body, however, the mechanism underlying brain lipid metabolism regulations remains a mystery. There is evidence to show that the brain’s energy metabolism largely depends on the temperature (*Guyton and Hall, 2006; Yu et al., 2012*). We therefore assume that cold modulates brain lipid metabolism in brain regions that express the genes encoding lipid metabolic enzymes sensitive to ambient temperature. Identifying the brain regions and genes modulated by ambient temperature is of significance in our understandings of the mechanisms of maintaining brain metabolism homeostasis.

Here, we combine cold exposure and brain region-selective genetic, molecular, and lipid metabolic assays, coupled with in vivo real-time two-color fiber photometry monitoring of lipid metabolic activity, to define cold-sensitive brain region(s) and lipid metabolism and to elucidate the involved mechanisms.

## Results

### Cold-induced brain region-selective gene expressions of lipid lipolytic markers

To investigate a potential for cold exposure to modulate gene expressions for lipid lipolysis and thermogenesis in the hypothalamus, mice were exposed to a cold (4 °C) chamber for 4 to 6 hours, a more physiologically relevant condition. Immediately after the cold challenge, mouse brains were acutely extracted and sectioned in ice-cold oxygenated artificial cerebrospinal fluids (ACSFs). Hypothalamic sections that respectively include the PVH, lateral hypothalamus (LH), dorsomedial hypothalamus (DMH), ventromedial hypothalamus (VMH), and arcuate nucleus (ARC) were transferred to ACSF-containing incubators. Micro-punches of these brain regions were subsequently made for gene assays of lipid metabolic markers.

To evaluate the effects of cold on the gene markers of lipolysis, we measured the mRNA levels of two key markers of lipolysis adipose triglyceride lipase (Atgl) and hormone sensitive lipase (Hsl). We observed that cold-challenged male mice showed a significant increase in the gene expressions of the both of Atgl (Fig. 1A_1_) and Hsl (Fig. 1A_2_) selectively in the PVH but not in other brain regions (Fig. 1B-E). We also assayed the gene expressions of thermogenic marker Ucp2 (Uncoupling protein 2) and additional thermogenic factors (Cidea, Prdm16). Cold did not significantly affect mRNA expressions of these thermogenic markers in all the examined regions in this study (Fig. 1). Cold did not significantly affect gene expressions of these lipolytic and thermogenic markers in female mice (Supplementary Fig. 1). These results suggest that cold exposure (4 ∼ 6 h) could induce a rapid lipolytic activity to release fatty acids primarily in the PVH in males. Because cold did not significantly affect lipolytic markers in other regions and did not affect thermogenic markers, we next focused on studying cold-induced lipid mobilization and lipolysis in the PVH in males.

**Figure 1.**
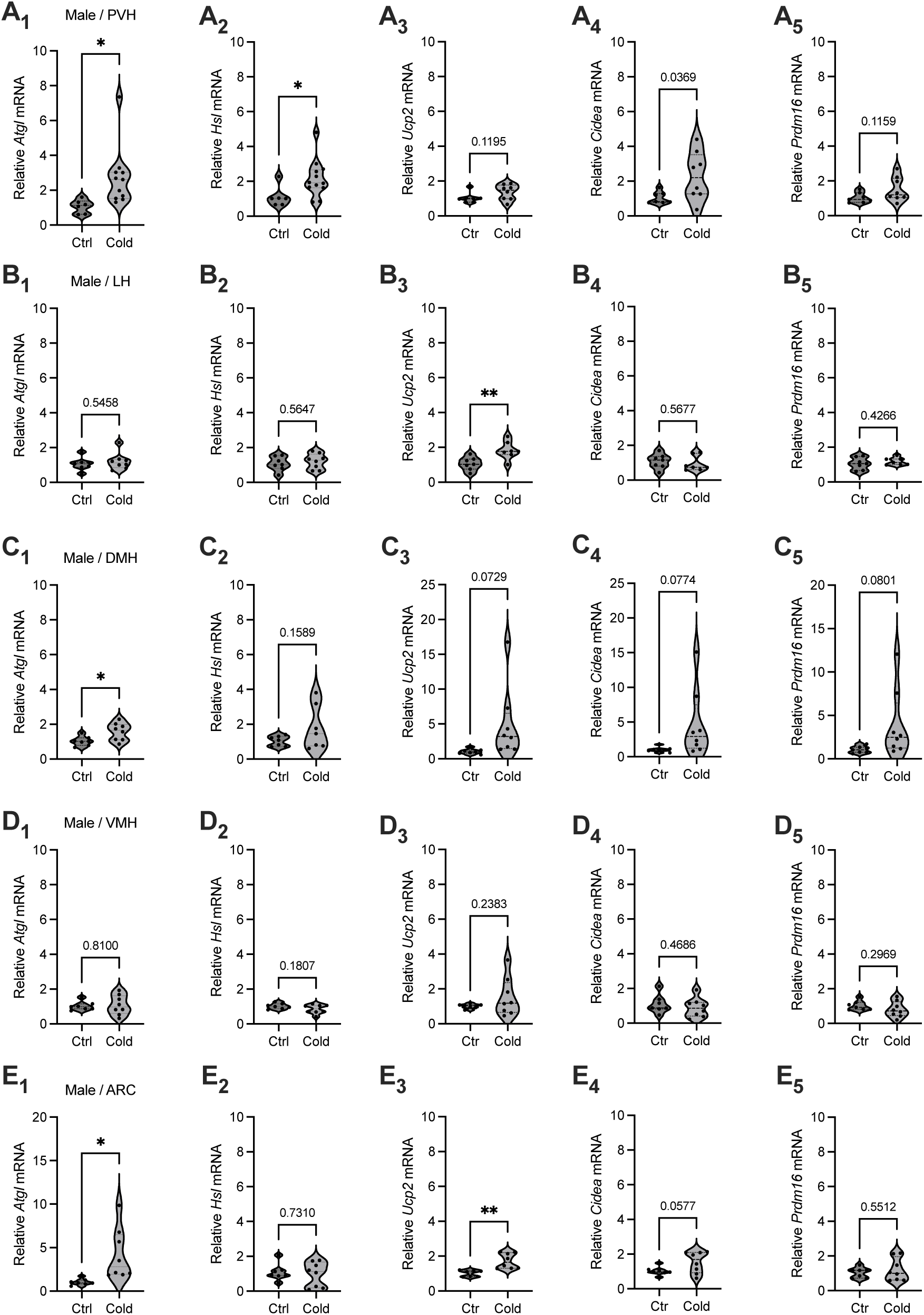
Brain region selective responses to cold. Micro-punches of PVH, LH, DMH, VMH, and ARC were made from male mice exposed to a cold chamber for 4-6 h, which were directly used for RT-qPCR of the gene markers of lipolysis and thermogenesis. (A_1_-A_5_) Group data of the lipolytic marker *ATGL* (A_1_) and *HSL* (A_2_) as well as thermogenic marker *Ucp2* (A_3_), *Cidea* (A_4_) and *Prdm16* (A_5_) in the PVH. (B_1_-B_5_) Group data of the lipolytic marker *ATGL* (B_1_) and *HSL* (B_2_) as well as thermogenic marker *Ucp2* (B_3_), *Cidea* (B_4_) and *Prdm16* (B_5_) in the LH. (C_1_-C_5_) Group data of the lipolytic marker *ATGL* (C_1_) and *HSL* (C_2_) as well as thermogenic marker *Ucp2* (C_3_), *Cidea* (C_4_) and *Prdm16* (C_5_) in the DMH. (D_1_-D_5_) Group data of the lipolytic marker *ATGL* (D_1_) and *HSL* (D_2_) as well as thermogenic marker *Ucp2* (D_3_), *Cidea* (D_4_) and *Prdm16* (D_5_) in the VMH. (E_1_-E_5_) Group data of the lipolytic marker *ATGL* (E_1_) and *HSL* (E_2_) as well as thermogenic marker *Ucp2* (E_3_), *Cidea* (E_4_) and *Prdm16* (E_5_) in the ARC. Data represent mean ± s.e.m. Student *t* tests were performed. \**p*<0.05. Each dot represents one animal in each group of all the panels.

**Figure 2.**
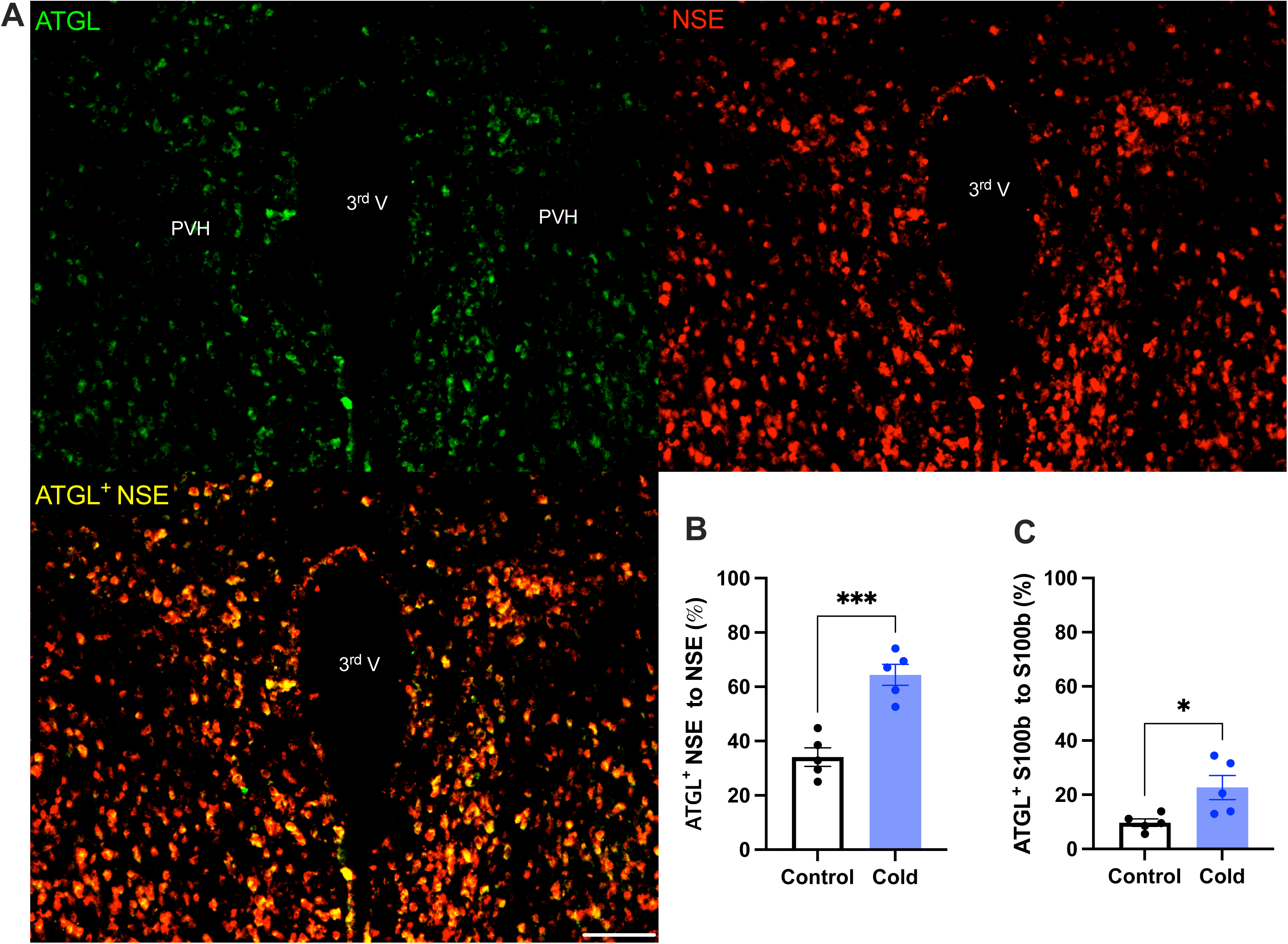
Cold increases protein expressions of ATGL in neurons and astrocytes in PVH. Control or cold (4-6)-challenged mice were perfused and fixed. Mouse brains were sectioned. ATGL, NSE, and S100b were stained using relevant antibodies respectively. (A) Representative images of ATGL (green), NSE (red), and ATGL/NSE overlay (yellow) signals in PVH sections from cold-challenged mouse. (B) Group data of relative ATGL/NSE overlay signals to total NSE signals in control and cold-challenged mice (n=5 each group). (C) Group data of relative ATGL/S100b overlay signals to total S100b signals in control and cold-challenged mice (n=5 each group). Data represent mean ± s.e.m. Student *t* tests were performed. \*\*\**p*<0.001, \**p*<0.05. Scale bar, 100 μm for (A). PVH, paraventricular of hypothalamus; 3^rd^ V, third ventricle.

### Cold increases the expressions of lipid lipolytic enzymes in the PVH

To verify the cold-induced effects on lipolytic gene expressions, we next stained the protein expressions of ATGL and HSL in PVH sections using antibodies against the ATGL and HSL respectively. Matching with the gene expression results, cold-challenged mice showed significant increases in ATGL (Fig. 2) and HSL (Fig. 3) expressions in both neurons and astrocytes respectively, compared to control mice.

**Figure 3.**
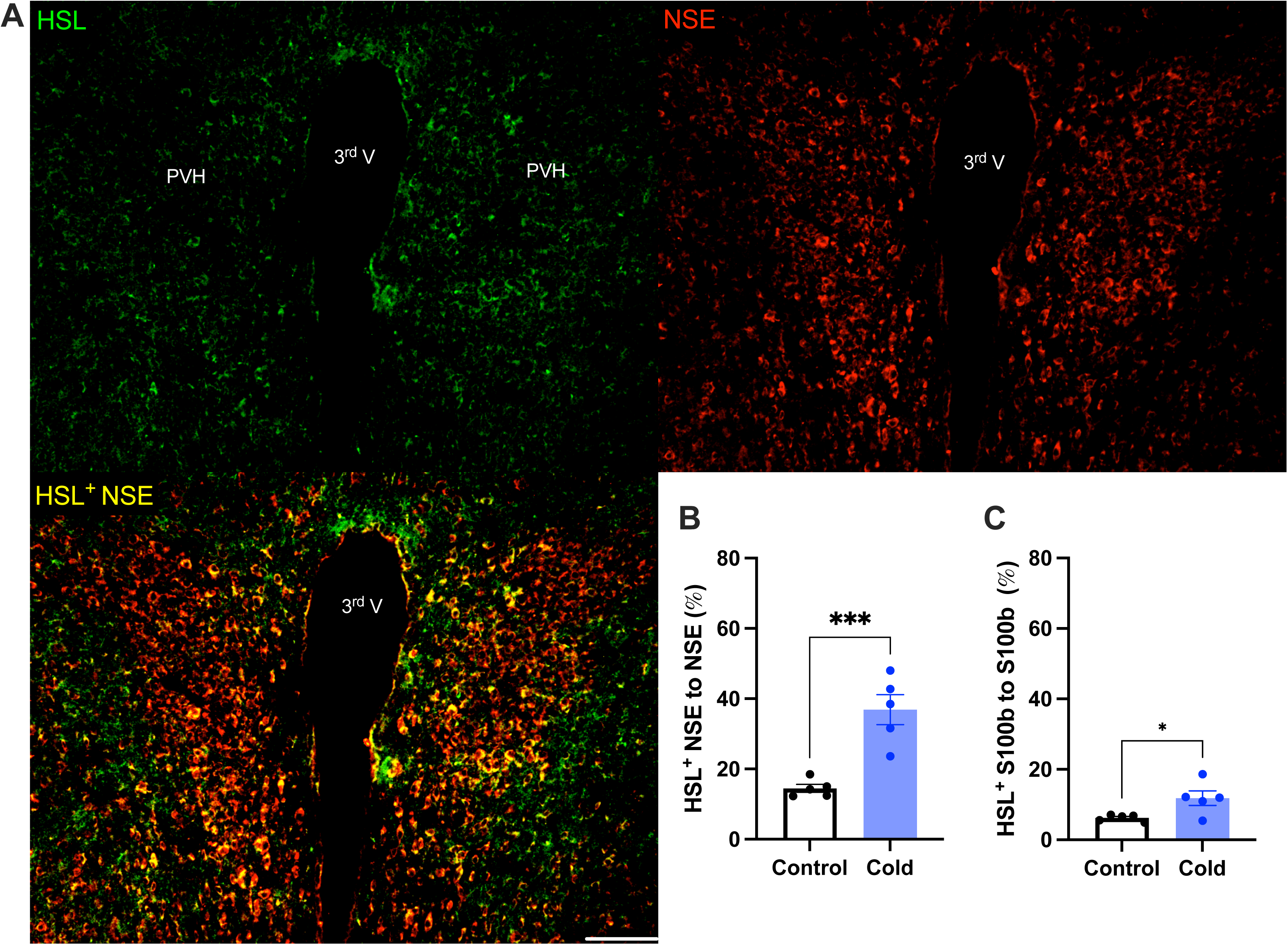
Cold increases HSL protein expressions in both neurons and astrocytes in PVH. Control or cold (4-6)-challenged mouse brains were sectioned. HSL, NSE, and S100b were respectively co-stained using relevant antibodies. (A) Representative images of HSL (green), NSE (red) and HSL/NSE overlay (yellow) signals in PVH sections from cold-challenged mouse. (B) Group data of relative HSL/NSE overlay signals to total NSE signals in control and cold-challenged mice (n=5 each group). (C) Group data of relative HSL/S100b overlay signals to total S100b signals in control and cold-challenged mice (n=5 each group). Data represent mean ± s.e.m. Student *t* tests were performed. \*\*\**p* < 0.001, \**p* < 0.05. Scale bar, 100 μm for (A) PVH, paraventricular of hypothalamus; 3^rd^ V, third ventricle.

Moreover, we examined whether and how cold modified the phosphorylation of the HSL, a key enzyme in the regulation of lipid lipolysis. After the cold exposure of cohort groups of mice, micro-punches of PVH sections were collected and directly used for western blots of phosphorylated HSL (p-HSL), HSL, and action. Cold significantly increased activation (S660) site phosphorylation of the HSL compared to controls (Fig. 4).

**Figure 4.**
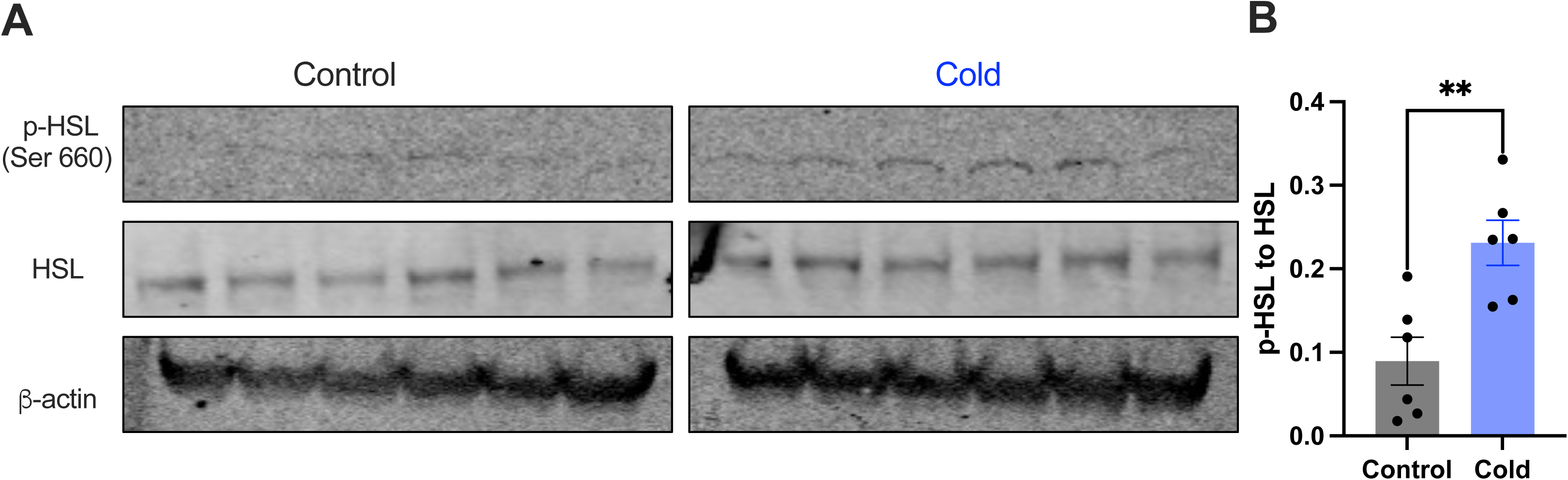
Cold increases the level of phosphorylated HSL in PVH. Micro-punches of PVH were collected in control and cold (4-6)-challenged mice, which were used for (A) western blots of p-HSL (Ser660), HSL, and β-action. (B) Group data of p-HSL fold change to HSL (n = 6 each group). Data represent mean ± s.e.m. Student *t* tests were performed. ** *p* < 0.01. Each dot represents one mouse.

### Cold-induced accumulation of lipid droplets (LDs)

It is increasingly appreciated that lipid droplets, endoplasmic reticulum (ER)-derived intracellular neutral lipid storage dynamic organelles (*Olzmann and Carvalho, 2019*), are predominantly accumulated in adipose tissues and liver under normal physiological conditions (*Farese and Walther, 2009; Murphy, 2001*). Also, there is evidence to indicate that glial cells such as astrocytes and tanycytes in the brain accumulate LDs under metabolic and hypoxic stress (*Geller et al., 2019; Smolic et al., 2021*). A recent elegant study shows that hyperactive neurons release fatty acids from phospholipids to formulate LDs in the brain and activation of neurons promotes lipolysis by increasing cytoplasmic lipases (*Ioannou et al, 2019*). These findings suggest that increased neuronal activity and accumulated LDs probably contribute to the cold-induced increase in the lipolytic markers we observed in this study.

To probe the mechanism of the upregulation of lipolytic markers in the PVH, we evaluated the ability for cold to accumulate LDs. After the cold challenge, mouse brains were perfused, fixed, and sectioned. Hypothalamic sections containing PVH were stained using the BODIPY 493 (BD493), a LD probe which has been well developed and widely applied to measure LD number and area in fixed tissues and live cells respectively (*Geller et al., 2019; Ioannou et al., 2019; Liu et al., 2015; Long et al., 2012; Smolic et al., 2020; Spangenburg et al., 2011*). Interestingly, we observed that a short-term (30 min ∼ 1 h) cold exposure induced an increase in the LD area in the PVH (Fig. 5A-C). We verified LDs by using the cytoplasmic LD-binding protein perilipin-2 (Fig. 5D-F). To define the cell type selectivity of LD accumulation induced by cold, we co-stained LDs with the BD493, astrocytes with S100b, and neurons with neuronal marker NSE respectively in PVH sections. We observed that cold exposure significantly increased the BD493-positive S100b-expressing astrocytes but not NSE-expressing neurons (Supplementary Fig. 2), consistent with the previous reports that LDs were primarily accumulated in glial cells (*Bailey et al., 2015; Liu et al., 2015; Marschallinger et al., 2020; Smolic et al., 2020*). However, a longer (4 ∼ 6 h) cold exposure reduced the number of LDs (Supplementary Fig. 3), which might be due to cold-induced liberations of fatty acids from the LDs to mitochondria for β-oxidation.

**Figure 5.**
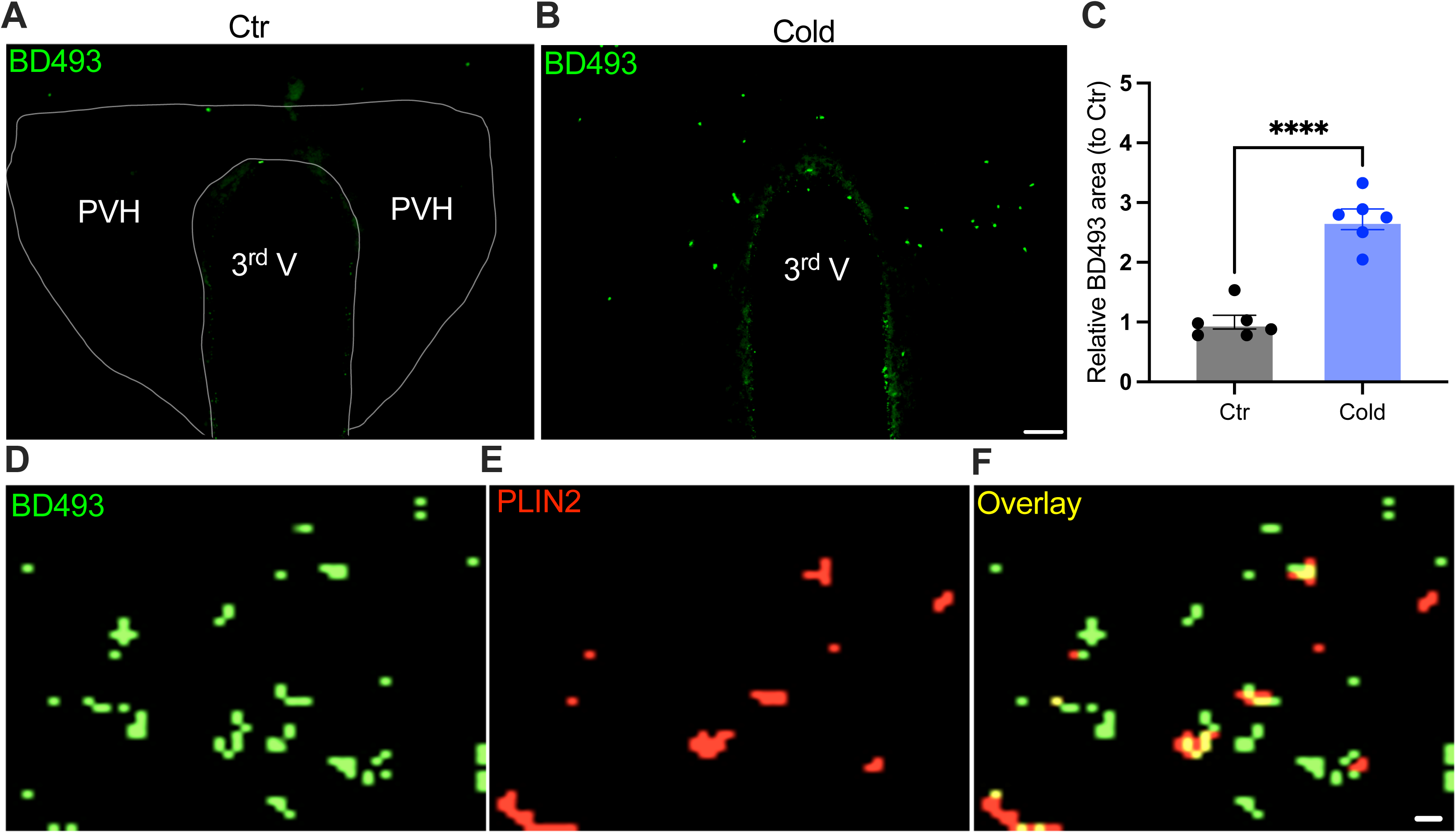
Cold-induced LD accumulation. Control or cold (30 min∼1 h)-challenged mice were perfused and fixed. Mouse brains were sectioned. LDs in the PVH were labelled using BODIPY493 (BD493). (A, B) Representative images of BD493 signals in PVH sections from one control (A) and one cold-challenged mouse (B). (C) Group data of relative BD493-labelled area in PVH in control and cold-challenged mice (n = 6 each group). (D-F) Sample images of BD493 (D) and perilipin2 (PLIN2) (E) and overlay (F) signals in the PVH. Data represent mean ± s.e.m. Student *t* tests were performed. \*\*\*\**p* < 0.0001. Scale bars, 50 μm for (A, B), and 1 μm for (D-F). PVH, paraventricular of hypothalamus; 3^rd^ V, third ventricle.

### Cold increases Fos expressions in PVH

We next examined whether and how cold modified cell activities within the PVH. Mice were placed in a cold chamber before mouse brains were perfused and fixed. PVH sections (40 μm in thickness) were stained using anti-Fos antibodies. Mice subjected to the cold challenge showed increased expressions of the activity indicator Fos in the PVH as compared to control mice (Fig. 6A-C).

**Figure 6.**
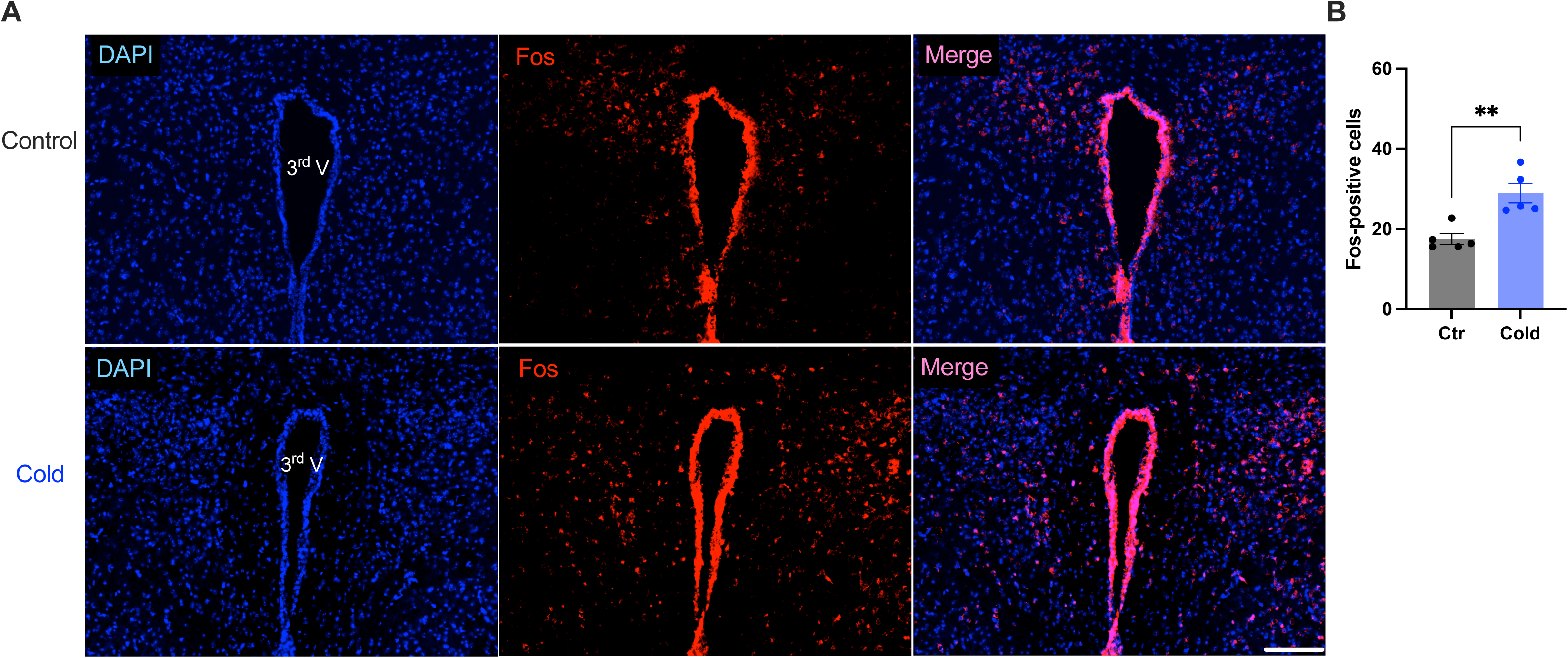
Cold increased Fos expressions in the PVH. Control or cold-challenged mouse brains were perfused and fixed and sectioned. Fos in the PVH were labelled using anti-Fos antibodies. (A) Representative images of Fos signals in PVH from one control (top) and one cold-challenged mouse (bottom). (B) Group data of Fos-positive cells in PVH in control and cold-challenged mice (n = 5 each group). Data represent mean ± s.e.m. Student *t* tests were performed. \*\**p*<0.01. Scale bars, 100 μm for (A). 3^rd^ V, third ventricle.

### *In vivo* real-time photometry monitoring of cold-induced lipid peroxidation

As lipid peroxidation is essential in neuronal activity-dependent accumulation of LDs (*Ionanna et al., 2019*), we next evaluated the capability for cold to induce lipid peroxidation. Fiber photometry has recently been applied to detect the levels of fluorescent biosensors *in vivo* by us (*Chen et al., 2022*) and others (*Andersen et al., 2023; Sun et al., 2018*). To achieve this goal, we took advantage of the BODIPY581/591 C11 (BD-C11) ratiometric lipid peroxidation sensor, coupled with *in vivo* time-lapse two-color photometry monitoring approach. We developed and validated this approach to simultaneously monitor both red and green signals in one brain region through a single implanted photometry fiber connected to a two-color photometry system, for the BD-C11 sensor as it shifts its fluorescence emission peak from 590 (red) and 510 (green) nm when oxidized. A custom-made <u>O</u>ptical fiber <u>m</u>ultiple <u>F</u>luid injection <u>C</u>annula (OmFC) implanted over the PVH was used for both BD-C11 injection and photometry monitoring of the two signals in freely behaving mice. Intra-PVH injection of the BD-C11 sensor through the OmFC was performed 4 h prior to placing the mice in a temperature-controlled chamber. Cold potently increased the BD-C11 ratio (green to red), indicating increased lipid peroxidation (Fig. 7A). To verify this result, we treated mice with a lipid soluble antioxidant α-tocopherol (α-TP) to inhibit lipid peroxidation. Compared to vehicle-treated mice (Fig. 7B), prior inhibition of lipid peroxidation with α-TP blunted the cold-induced effects on lipid peroxidation (Fig. 7C). These results demonstrate that cold induces lipid peroxidation in the PVH and that our combined photometry and BD-C11 approach can be reliably applied to evaluate the dynamics of brain lipid metabolism in a spatiotemporal manner.

**Figure 7.**
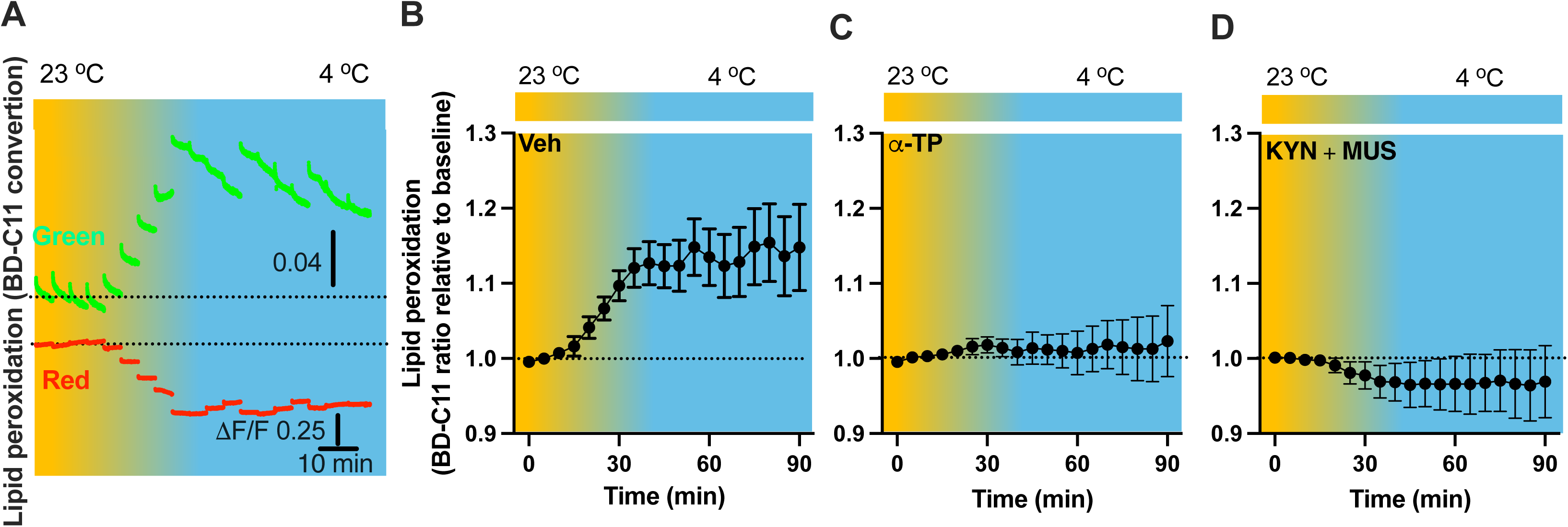
Cold-induced cell activity-dependent lipid peroxidation in the PVH. Mice were injected with the lipid peroxidation indicator BD-C11 in the PVH via the implanted OmFC cannula 4 h before placing them in a temperature-controlled chamber. Two-color fiber photometry was applied for time-lapse monitoring of BD-C11 convertion via the optic fiber of the OmFC. (A) Representative traces of real-time photometry monitoring of red and green signals simultaneously. (B-D) Group data of relative BD-C11 ratio to baseline in mice treated with vehicle (B, n=6), α-TP (C, n= 4), and KYN+MUS (D, n=4). Data represent mean ± s.e.m. Student *t* tests were performed.

### Lipid peroxidation is under the control of cell activity

We next sought to examine whether lipid peroxidation is under the control of cell activity. To examine whether cold alters cell activity, we transduced PVH cells with the Ca^2+^ indicator GCaMP_6f_- and placed the PVH transduced and photometry fiber implanted mice in a cold chamber (Supplementary Fig. 4A). Cold exposure increased the intensity of PVH GCaMP_6f_ signals (Supplementary Fig. 4 B and C), consistent with the increased Fos expressions (Fig. 6). To test inhibit cell activity, we utilized the GABA_A_ receptor agonist muscimol (MUS) (*Barbalho et al., 2009; Sanders and Shekhar, 1995*) and glutamate receptor antagonist Kynurenic acid (KYN) (*Yoshida et al., 2012*). To verify the inhibitory effect of the combined two chemicals, we performed intra-PVH injection of MUS and KYN in PVH GCaMP_6f_ – transduced and fiber implanted mice. Thirty min post the MUS and KYN administration via intra-PVH injection was able to reduce cell activity, as indicated by the decreased intensity of GCaMP_6f_ signals, compared to ventricle treatment (Supplementary Fig. 4D and E).

To test whether cell inactivation could diminish the cold-induced effect on lipid peroxidation, intra-PVH injections of MUS and KYN were performed 30 min before placing the previously BD-C11 injected mice in a cold chamber and performing photometry monitoring of BD-C11 signals. Administration of the MUS and KYN prevented the cold-induced increase in the BD-C11 ratio (Fig. 7D), suggesting that cell activation is required for the cold-induced lipid peroxidation.

### Cell inactivation prevents cold-induced lipolysis

To evaluate lipolysis in the PVH, we performed intra-PVH injections of the EnzCheck lipase substrate through an implanted OmFC. The lipase substrate shifts its fluorescence to emission peak 515 nm (green) in the presence of lipases, and we detected the green signals using our photometry system in a temporal manner. Intra-PVH injection of EnzCheck lipase substrate was performed 4 h before placing mice in a temperature-controlled chamber. Cold induced a potent increase in the green signals generated from the lipase substrate, indicating increased lipolysis (Fig. 8A-B). To verify this, we injected mice with the pan-lipase inhibitor diethylumbelliferyl phosphate (DEUP) via intra-PVT injections to inhibit lipolysis. Cold-induced lipolysis was reduced with the treatment of DEUP compared to vehicle-treated mice (Fig. 8C). These results are consistent with the cold-induced increase in lipolytic markers (Figs. 1 - 4), further indicating that cold induces lipolysis in the PVH by increasing cytoplastic lipases. To examine whether cell activity is required for the cold-induced lipolysis, in cohort groups of PVH-implanted mice which were treated with the lipase substrate via intra-PVH injections, we performed intra-PVH injections of the MUS and KYN 30 min before placing the mice in a cold chamber. Administration of MUS and KYN prevented the cold-induced conversion of the lipase substrate, indicating decreased lipolytic activity (Fig. 8D). These findings suggest that cold induces a rapid activity-dependent lipid lipolytic effect.

**Figure 8.**
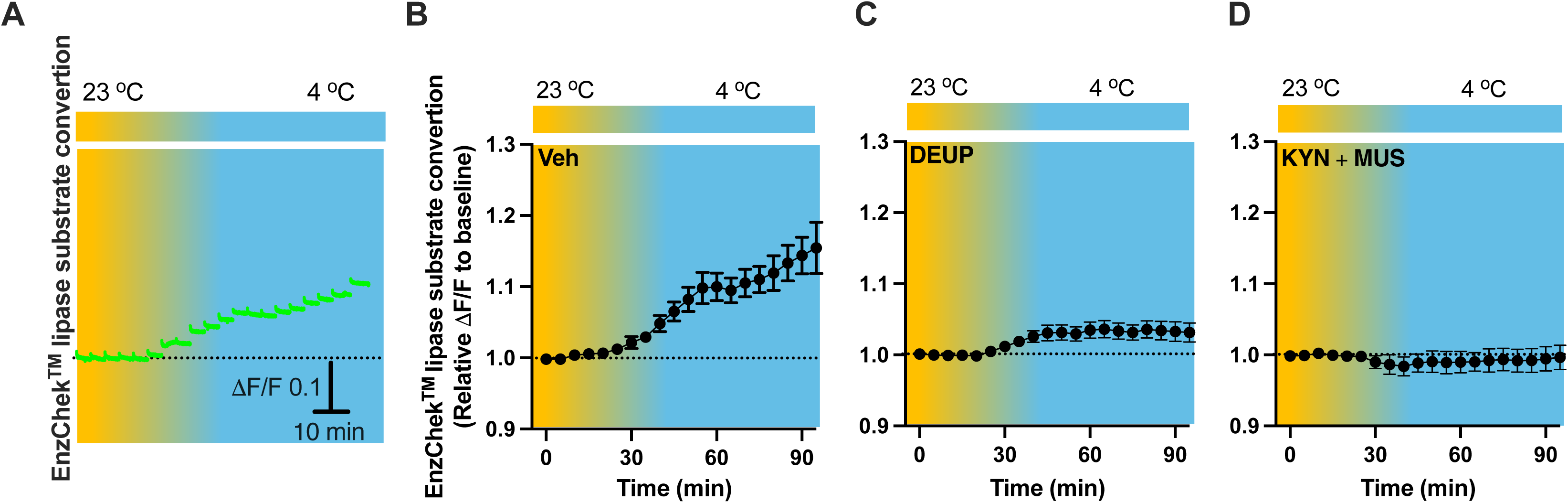
Cold-induced cell activity-dependent lipid lipolysis. Mice were injected with the lipase substrate in the PVH via the implanted OmFC cannula 4 h before placing them in a temperature-controlled chamber. Two-color fiber photometry was used for time-lapse monitoring of lipase substrate convertion via the optic fiber of the OmFC. (A) Representative traces of real-time photometry monitoring of green signals. (B-D) Group data of lipase substrate convertion in mice treated with vehicle (B, n=4), α-TP (C, n=4), and KYN+MUS (D, n=4). Data represent mean ± s.e.m. Student *t* tests were performed.

### Cell inactivation reduces cold-induced LD accumulation

To verify our results of LDs detected in the fixed tissues (Fig. 5), we next performed time-lapse photometry monitoring of dynamics of LDs *in vivo* in freely behaving mice. To achieve this goal, we implanted an OmFC cannula in PVH for LD marker BD493 (green) injections and monitoring in mice. Matching with the results collected from the fixed sections, cold induced an increase in the intensity of BD493 signals (Fig. 9A and B). This result indicates that cold increases the accumulation of LDs which could be directly used for lipolysis by releasing fatty acids to mitochondria for β-oxidation. The lipid soluble antioxidant α-TP blocked the cold-induced LD accumulation (Fig. 9 C). To define a role of cell activity in modulating LDs, we performed intra-cannula injections of both MUS and KYN in BD493-injected mice in the PVH. MUS and KYN administration blunted the cold-induced accumulation of LDs, as cold failed to increase the BD493 signals in MUS and KYN treated mice compared to controls (Fig. 9D).

**Figure 9.**
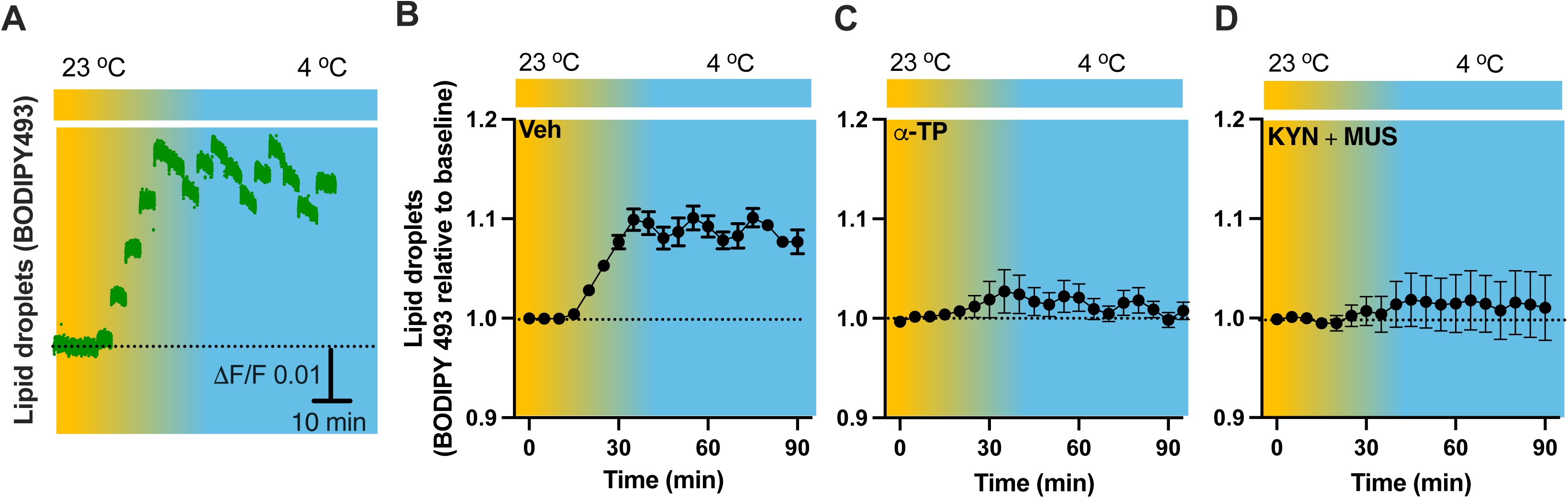
Cold-induced cell activity-dependent LD accumulation. Mice were injected with the LD marker BODIPY493 in the PVH via the OmFC cannula 4 h before placing them in a temperature-controlled chamber. Fiber photometry was used for time-lapse monitoring of BODIPY493 signals via the optic fiber of the OmFC. (A) Representative traces of real-time photometry monitoring of BODIPY493 signals. (B-D) Group data of BODIPY493 in mice treated with vehicle (B, n=4), α-TP (C, n= 4), and KYN+MUS (D, n=6). Data represent mean ± s.e.m. Student *t* tests were performed.

## Discussion

For survival, it is crucial for individuals to precisely modulate lipid metabolism in both the peripheral tissues and brain in mammalian animals. Extensively studies have been focused on lipid metabolism in peripheral tissues particularly adipose tissues, while little or nothing was known about that in the brain. This was probably in part because of limited techniques. By taking advantage of combined coordinated neurobiology methods and lipid metabolic assays, we provided in this study the *in vivo* evidence that cold is capable to induce acute cell activity-dependent central nervous system lipid metabolic activities, including lipid peroxidation and mobilization, lipid droplet formation, and lipid lipolysis in the hypothalamus. These results fill in a missing but important gap in our understanding of brain region selective lipid metabolism in physiological conditions such as cold applied in this study. Also, our *in vivo* time-lapse fiber photometry approach detecting probe-labelled lipid metabolites would provide an alternative method to evaluate the dynamics of lipid metabolism in live animals. We believe our studies will also be paradigmatic for understanding abnormal lipid metabolism in brain regions modifying energy metabolism in metabolic disorders such as obesity.

The third ventricle adjacent PVH is a conserved brain region (*Machluf et al., 2011*), composed of different cell populations including those multiple brain regions-projecting parvocellular neurons and magnocellular neurons. Particularly, PVH plays varied crucial roles in modulating the hypothalamic-pituitary-adrenal (HPA) axis and the hypothalamic-pituitary-thyroid axis (HPT) (*Swanson et al., 1983*). This determines a key role of the PVH in modulating body functions, energy metabolism and glucose homeostasis in health and disease. Our findings in this study show that cold increases the expressions levels of Fos, lipid peroxidation, LD accumulation, and lipolytic markers in the PVH. These findings reveal a new role for the PVH in preventing fatty acid toxicity in pathological conditions.

Activated neurons would induce phospholipid peroxidation to release toxic fatty acids. One recent elegant study (*Ioannou et al., 2019*) shows that nearby astrocytes have the capacity to endocytose neuron-released fatty acids (FAs), store them in LDs, and liberate FAs from LDs through lipolytic processes to mitochondria for β-oxidation. Consistently, our data in this study demonstrate that cold-induced lipid peroxidation and lipolysis in the brain are also under the control of cell activities. Our data also show that cold induces increased expressions of ATGL and HSL in both neurons and astrocytes (Figs. 2 and 3). These findings suggest that both neurons and astrocytes play important roles in inducing lipid metabolism and fatty acid accumulations in the brain in certain physiological and pathological conditions.

Lastly, the brain is highly thermal sensitive (*Brooks, 1983*). Identifying brain regions sensitive to ambient temperature is significant and important in our understandings of brain energy metabolism and homeostasis, as which might provide potential targetable sites in the treatment of brain lipid metabolism-associated neurological disorders.

## Methods

Experimental protocols were approved by the Institutional Animal Care and Use Committees at the Albert Einstein College and conducted following the U.S. National Institutes of Health guidelines for animal research.

### Animals

C57/BL6J wild type mice (#000664, Jackson Laboratory) have been described previously and are available from The Jackson Laboratory. Both male and female mice (age 8-12 weeks) were used at the start of experiments. Mice were group-housed 3–5 mice per cage in humidity- and temperature (22–25 °C)-controlled rooms on a 12-hour light:dark cycle (lights on from 8:00 a.m. to 8:00 p.m.) with *ad libitum* access to water and mouse regular chow (#5001, LabDiet). Mice were single-caged after they received viral transductions with or without guide or optic fiber cannula insertion until all the experimental procedures were finished.

### Pharmacology

All the chemicals were purchased from Sigma except where noted. For the experiments requiring intra-PVH injections, an injector with 1-mm extension beyond the custom-made optical fiber multiple fluid injection cannula (OmFC) (Doric Lens) implanted over the PVH was attached by polyethylene tubing to a Hamilton syringe. The injection was performed at a speed of 50 nl per min for 4 min using a matched fluid injector consisiting of a 1.25 mm ferrule and a sleeve connector, and the injector was withdrawn 10 min after the final injection. Grip cement (DENTSPLY) was used to anchor the cannula to the skull, and a plug was inserted to keep the cannula from becoming clogged when the injector was not in place. Mice were then returned to the home cage for one week at least before the experiments. The amount for intra-PVH injection was 200 nl of vehicle or chemicals (in μM): 100 BODIPY™ 493/503 (D3922, Invitrogen), 100 BODIPY™ 581/591 C11 (D3861, Invitrogen), 1 MitoSOX™ Mitochondrial Superoxide Indicators (M36005, Invitrogen), 1 EnzChek™ Lipase Substrate 505/515 (E33955, Invitrogen), 100 Diethylumbelliferyl phosphate (D7692, Sigma), and 500 α-Tocopherol (#258024, Sigma); or 200 nl of 250 pmol Muscimol (M1523, Sigma) and 100 mol Kynurenic acid (K3375, Sigma).

### Stereotaxic OmFC implantations for intra-PVH injections and photometry

Following our previously documented protocols (*Chen et al, 2022; Zhang et al., 2020*), mice were anesthetized with 3% isoflurane to induce the anesthesia and with 1.5–2.0% isoflurane to maintain anesthesia during the surgery and placed in a stereotaxic frame (Kopf Instruments). A small incision was made in the skin of the head, a small hole was drilled on the skull, and mice were implanted with a custome-made optical fiber multiple fluid injection cannula (OmFC) (Doric Lens) over the PVH (coordinates from bregma: AP −0.8 mm, 0.2 mm from midline, DV −4.0 mm). The cannula was fixed to the skull with stainless steel screws and dental cement. After the surgery, all mice received meloxicam (5 mg/kg) and continued to be housed individually. Two weeks after surgery, the mice were briefly anesthetized and inserted with the fluid injector (with 1 mm protrusion) into PVH. For intra-PVH injection, the injector was attached to a Hamilton syringe through a polyethylene tube and the mice received 200 nl of vehicle, chemicals as listed in the Pharmacology section, or viral vectors (AAV_5_-CamKII-GCaMP_6f_-WPRE-SV40, addgene#100834, titer, 2.3 × 10^13^ GC/ml) or chemicals as stated above, at a rate of 50 nl/min.

### Micro-punches of hypothalamic nuclei

As described in our previously published studies (*Qi and Yang, 2015; Chen et al, 2022; Zhang et al., 2020*), acute brain slices that include hypothalamic PVH, LH, DMH, VMH, and ARC respectively were prepared. Briefly, mice were first placed in a cold chamber (4 °C) for 30 min or 4-6 h respectively, as described in the text and figure legends. After the cold exposure, mice were deeply anesthetized with isoflurane and decapitated. The mouse brains were dissected rapidly and placed in ice-cold oxygenated (95% O_2_ and 5% CO_2_) solution containing the following (in mM): 110 choline chloride, 26.2 D-glucose, 2.5 KCl, 1.25 NaH_2_PO_4_, 2 CaCl_2_, 7 MgSO_4_, 11.6 Na-L-ascorbic acid, and 3.1 Na-pyruvate. Coronal brain slices (280 μm thick) were cut with a vibratome (Leica; VT 1200 S) and maintained in an incubation chamber containing artificial cerebrospinal fluids (aCSFs) (in mM): 119 NaCl, 25 NaHCO_3_, 11 D-glucose, 2.5 KCl, 2.5 CaCl_2_, 1.3 MgSO_4_, and 1 NaH_2_PO_4_. Micro-punches of the PVH, LH, DMH, VMH and ARC were performed using a punch (1.5 mm diameter; Stoelting#57403) or a pipette tip (1.5 mm diameter). Six micro-punches of each region were obtained from each mouse were respectively collected and immediately placed in 200 μl Trizol reagent (Invitrogen) and stored at −80 °C until analysis.

### Total RNA extractions and Real-time qPCR (RT-qPCR)

Total RNA was extracted using Trizol reagent (Invitrogen) according to manufacturer’s instruction. Briefly, the collected tissues were lysed in 200 μl of Trizol reagent (Invitrogen) and 40 μl of chloroform was added into each tube and subsequently vortexed for 15 s. After incubation for 2 min, centrifuged at 12,000 x g for 10 min (4 °C). The aqueous phase was transferred to fresh tube and equal volume of isopropanol was added. The tube was incubated for 20 min and centrifuged at 12,000 × g for 10 min (4 °C). The RNA pellet was washed with 70% ethanol by vortex and centrifuged at 12,000 x g for 10 min (4 °C). cDNA was synthesized by High-Capacity cDNA Reverse Transcription Kit (Applied Biosystems) following the manufacturer’s instructions. Real-time qPCR was performed using QuantStudio 3 instruments (Applied Biosystems). The mixture for qPCR reaction was prepared in a final volume of 20 μl containing 1 μl cDNAs and 10 μl of LightCycler 480 SYBR Green I Master (Roche) in the presence of primers at 500 nM. The specific primer sequences used were the following:

The specific primer sequences used were the following:

ATGL forward, 5′-CCAACACCAGCATCCAGT-3′;

ATGL reverse, 5′-CAGCGGCAGAGTATAGGG-3′;

HSL forward, 5′-CGCCATAGACCCAGAGTT-3′

HSL reverse, 5′-TCCCGTAGGTCATAGGAGAT-3′;

Cidea forward, 5′-TGCTCTTCTGTATCGCCCAGT-3′

Cidea reverse, 5′-GCCGTGTTAAGGAATCTGCTG-3′;

Prdm16 forward, 5′-CCACCAGCGAGGACTTCAC-3′

Prdm16 reverse, 5′-GGAGGACTCTCGTAGCTCGAA-3′;

Ucp2 forward, 5′-CAGAGCACTGTCGAAGCCTA-3′

Ucp2 reverse, 5′-GTATCTTTGATGAGGTCATA-3′;

Actb forward, 5′-GCTGTCCCTGTATGCCTCT-3′

Actb reverse, 5′-GTCTTTACGGATGTCAACG-3′;.

The relative level of expression was calculated using the comparative 2^-ΔΔCT^ method.

### Two-color two-channel fiber photometry (FP)

Briefly, three excitation wavelengths were used: 560, 505 / 490, and 405 nm. Excitation lights were generated through fiber-coupled two connectorized LEDs (CLED_560 for 560 nm; CLED_505 / 490 for 505 or 490 nm; CLE_405 for 405 nm; Doric Lenses) driven by a four-channel LED driver (LEDD_4; Doric Lenses). The LEDD_4 was controlled by a fiber photometry console (FPC; Doric Lenses) connected to a computer. Excitation lights were passed through two fluorescence MiniCubes (iFMC6_IE(400-410)_E1(460-490)_F1(500-540)_E2(555-570)_F2(580-680)_S)-6 ports with 2 integrated photodetector heads. The single detector measures both signals within the fluorescence detection windows from 500-540 nm and 580-680 nm band. The combined excitation light was sent into a patch cord made of a 400 µm core, 0.48 NA, low-fluorescence optical fiber (Doric Lenses). The patch cord was connected to the implanted OmFC consisting of a 1.25 mm diameter optic fiber via a sleeve (Doric Lenses; Zirconia Sleeve 1.25 mm with black cover; Sleeve_ZR_1.25-BK). Both green and red fluorescence signals were collected through the same patch cord and passed through the same Minicube and focused onto a Fluorescence Detector Head (FDH; Doric Lenses). The photometry experiments were run in a Lock-in mode, and the acquisition rate was set to 12.0 ksps*C controlled by Doric Neuroscience Studio software (Doric Lenses). The FP experiments were performed 5 min after connecting the optic fibers to the animals. We processed the signals using the Doric Neuroscience Studio software (V5.3.3.14) to calculate the normalized fluorescence variation of the images (ΔF/F), and averaged the signals at 1-s bins. To avoid or minimize bleaching over time, we performed patch cord photobleaching for 12 h before each experiment, reduced the illumination power outputs as much as possible, and recorded the signals for 30 s every 5 min. We used a rotary joint for long-term photometry recordings in freely moving animals.

### Brain histology and immunofluorescence

Mice were euthanized and transcardially perfused with 1 x phosphate buffered saline (PBS, pH 7.4) followed by 4% paraformaldehyde in phosphate buffer (PFA, pH 7.2). Mouse brains were removed and post-fixed in 4% PFA overnight. Fixed brains were transferred to 30% sucrose in PBS for cryoprotection. Next, 30 μm coronal sections were cut in a freezing cryostat (Leica). For immunofluorescence, slices were washed three times in 1 x PBS for 10 min each and heat-incubated with target antigen retrieval solution (Invitrogen) for 2 min at 95 °C. And then, the slices were washed three times in 1 x PBS for 10 min each, followed by permeabilization in 1% triton X-100 solution in 1 x PBS for 40 min at room temperature. After blocking for 1 hour at room temperature, the slices were incubated with a mouse Fos antibody (1:150, sc-166940 AF647, Santa Cruz Biotechnologies), rabbit ATGL (1:150, bs-3831R-BF488, Bioss), rabbit HSL (1:150, bs-3223R-BF488, Bioss), rabbit NSE (1:200, bs-10445R-BF594, Bioss), and mouse S100b (1:200, bsm-10832-BF594, Bioss), respectively, for overnight at 4 °C in the dark. The slices were then rinsed three times in 0.1% triton X-100 solution in 1 x PBS for 10 min each. For lipid droplet staining, slices were washed three times in 1 x PBS for 10 min each and then incubated with 1% triton X-100 solution in 1 x PBS for 40 min. After blocking for 1 hour, the slices were stained with 20 μg/mL BODIPY 493/503 (Invitrogen, D3922) for 4 hours at room temperature in the dark. After the incubation, the slices were rinsed three times in 0.1% triton X-100 solution in 1 x PBS for 10 min each. For BODIPY 493/503 co-staining with perilipin-2, NSE, and S100b respectively, after permeabilization in 1% triton X-100 solution in 1 x PBS for 40 min and blocking for 1 hour at room temperature, the slices were incubated with rabbit perilipin-2 (1:200, CL594-15294, Proteintech), rabbit NSE, and mouse S100b antibodies, respectively, for overnight at 4 °C in the dark. The slices were then washed three times in 0.1% triton X-100 solution in 1x PBS for 10 min and incubated with 20 μg/mL BODIPY 493/503 (Invitrogen, D3922) for 4 hours at room temperature in the dark and then washed three times in 0.1% triton X-100 solution in 1x PBS for 10 min each time. The stained slices were dried and mounted with mounting medium (0100–20, Southern Biotech). Images were taken using the All-in-One Fluorescence Microscope (BZ-X800E Keyence) and analyzed using the BZ-X800 Analyzer (Keyence).

### PVH tissue western blot

Acute coronal brain slices (260-μm thickness) were collected following the procedures as stated in the above. Six micro-punches of PVH sections from each mouse were made and immediately placed in 40 μl cold lysis buffer containing 50 mM Tris-HCl, 150 mM NaCl, 0.25% Na-deoxycholate, 1 mM EDTA, 1% NP-40, 1 mM PMSF (20 mM PMSF in 100% ethanol), and 1 protease inhibitor tablet (A32955, Thermo Scientific). The samples were homogenized in lysis buffer and centrifuged at 14,000 × rpm at 4 °C for 20 min. The supernatants were transferred to a fresh tube and used for Bradford assay to quantify the protein concentration.

Equal amounts of protein from each sample were denatured and boiled at 100 °C for 5 min. Proteins were separated by electrophoresis in 10% SDS-PAGE gel and transferred to PVDF membrane. The membrane was blocked with Protein-Free blocking buffer (927-80001, LI-COR) for 1 h and incubated overnight with relevant primary antibodies at 4 °C, including the p-HSL (1:1,000, PA5-64494, Invitrogen), HSL (1:1,000, PA5-17196, Invitrogen), and β-actin (1:5,000, MAB8929, R&D Systems). After washing in TBST, the membranes were incubated with secondary antibodies for 1 hour and a subsequent TBST washing. Images were acquired using Odyssey Classic Imager (LI-COR). The band intensity of HSL and p-HSL was quantified using the ImageJ.

## Author contributions

H.Y.M. and Y.Y. conceived, designed, and performed the experiments, analyzed data, and wrote the manuscript. Y.Y.Y. edited the manuscript.

## Acknowledgements

We thank all the members of the Yang laboratory for discussion and critical comments on this study. We thank the Doric Lenses for helping us build and adjust the two-channel two-color photometry rig. For the genetically encoded calcium indicator GCaMP_6f_ plasmids, we thank Dr. James M Wilson for depositing the plasmid of pENN-AAV-CamKII-GCaMP_6f_-WPRE-SV40. This work was supported by the NIH (R01 DK112759, R01 DK135717 to Y.Y.) and Einstein Research Foundation.

## Data and materials availability

All data are available in the main text or the supplementary materials.

