## Supplemental Info for "Cold induces brain region-selective cell activity-dependent lipid metabolism"

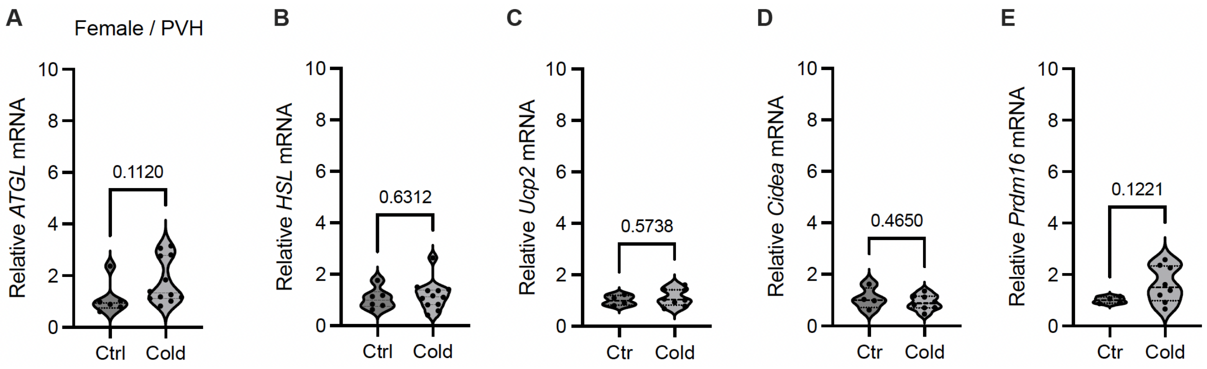


**Supplementary Figure 1.** Cold did not affect gene markers in PVH in female mice. Micro-punches of PVH were made from female mice exposed to a cold chamber for 4-6 h for RT-qPCR of the gene markers of lipolysis and thermogenesis. Group data of the lipolytic marker *ATGL* (A) and *HSL* (B) as well as thermogenic marker *Ucp2* (C), *Cidea* (D) and *Prdm16* (E) in the PVH. Data represent mean ± s.e.m. Student *t* tests were performed. Each dot represents one animal in each group of all the panels.

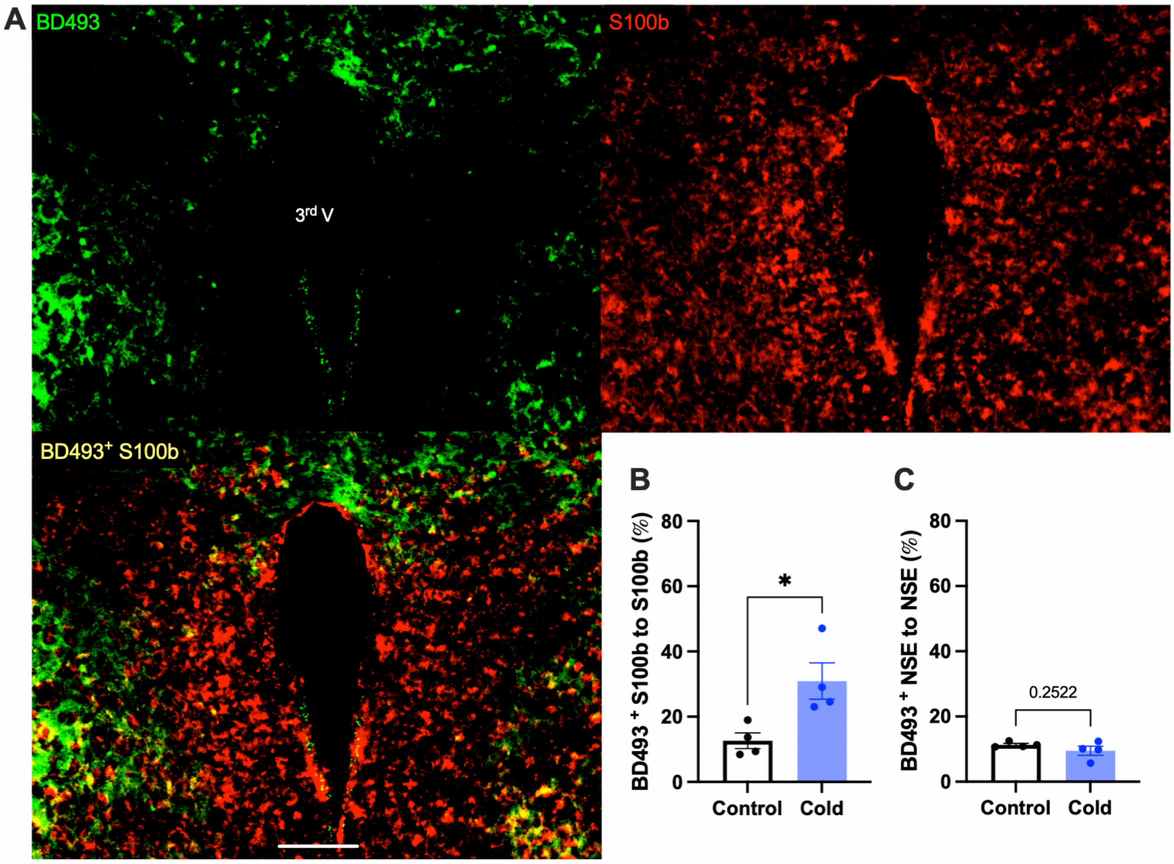


**Supplementary Figure 2.** Cold increases LD accumulation in astrocytes but not neurons. Control or cold (4- 6)-challenged mouse brains were sectioned. BD493, S100b, and NSE were respectively co-stained using relevant antibodies. (A) Representative images of BD493 (green), S100b (red), and BD493/S100b overlay (yellow) signals in PVH sections from cold-challenged mouse. (B) Group data of relative BD493/S100b overlay signals to total S100b signals in control and cold-challenged mice (n = 4 each group). (C) Group data of relative BD493/NSE overlay signals to total NSE signals in control and cold-challenged mice (n = 4 each group). Data represent mean ± s.e.m. Student *t* tests were performed. *$p$<0.05. Scale bar, 100 μm for (A). 3rd V, third ventricle.

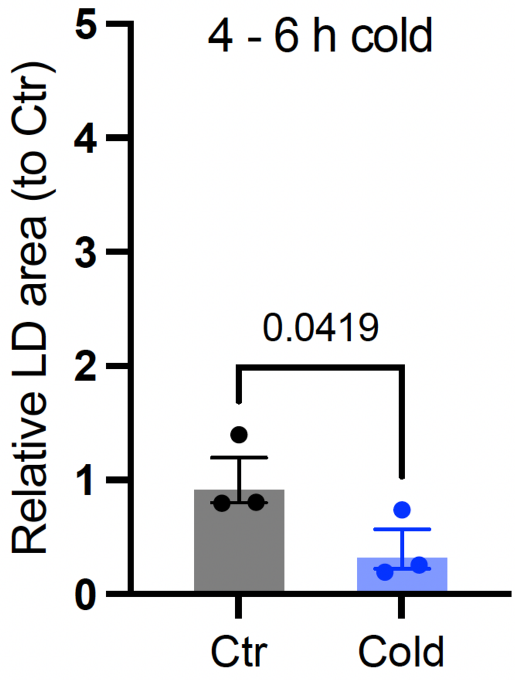


**Supplementary Figure 3.** Group data of relative BD493-labelled area in PVH in control or cold-challenged (4 ~ 6 h) mice (n=3 each group). Data represent mean ± s.e.m. Student *t* tests were performed.

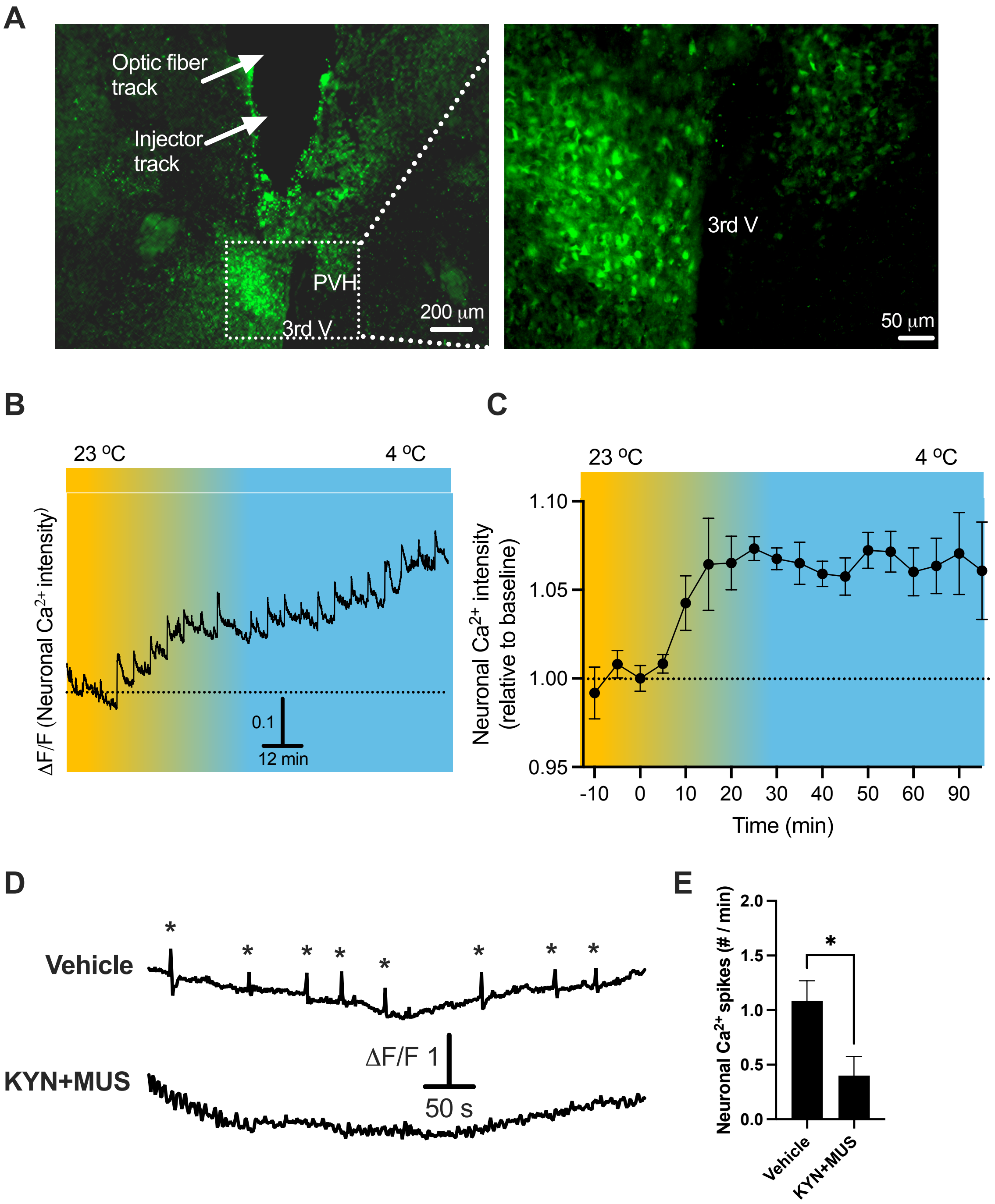


**Supplementary Figure 4.** Neuronal inhibition by KYN+MUS. (A) Representative images of GCaMP6f expressions in PVH neurons and cannula track. (B) Representative trace of neuronal GCaMP6f signals recorded in a time lapse manner in a cold chamber. (C) Group data of (B, n=3). (D) Representative traces of neuronal GCaMP6f signals recorded from vehicle (top) and KYN+MUS (bottom)-injected mice, and stars indicate $Ca^{2+}$ spikes. (E) Group data of relative GCaMP6f-labelled $Ca^{2+}$ spikes in PVH neurons in vehicle or KYN+MUS-injected mice (n=3 each group). Data represent mean ± s.e.m. Student *t* tests were performed. **p*<0.05 for (E).
